# Gut microbial ecosystems differ across metabolic and obesity phenotypes

**DOI:** 10.1101/2025.04.25.650405

**Authors:** Blanca Lacruz-Pleguezuelos, Guadalupe X Bazán, Sergio Romero-Tapiador, Gala Freixer, Jorge Fernández-Cabezas, Elena Aguilar-Aguilar, Adrián Martín-Segura, Lara P. Fernández, Isabel Espinosa-Salinas, Ana Ramírez de Molina, Aythami Morales, Ruben Tolosana, Javier Ortega-Garcia, Vera Pancaldi, Laura Judith Marcos-Zambrano, Enrique Carrillo de Santa Pau

**Affiliations:** Computational Biology Group. IMDEA Food, CEI UAM+CSIC. Carretera de Cantoblanco, 8. 28049 Madrid (Spain); UAM Doctoral School, Universidad Autónoma de Madrid, Spain; GENYAL Platform. IMDEA Food, CEI UAM+CSIC. Carretera de Cantoblanco, 8. 28049 Madrid (Spain); Biometrics and Data Pattern Analytics Lab, Escuela Politécnica Superior, Universidad Autónoma de Madrid, Spain; Department of Pharmacy and Nutrition, Faculty of Biomedical and Health Sciences Universidad Europea de Madrid, Calle Tajo s/n, Villaviciosa de Odon, 28670, Spain; Molecular Oncology Group. IMDEA Food, CEI UAM+CSIC. Carretera de Cantoblanco, 8. 28049 Madrid (Spain); Centre de Recherches en Cancérologie de Toulouse, CRCT, Université de Toulouse, Inserm, CNRS, Toulouse, France

**Author notes:** Address correspondence to Laura Judith Marcos-Zambrano,; Enrique Carrillo de Santa Pau.

## Abstract

Obesity is a heterogeneous condition comprising a continuum of phenotypes with various metabolic and inflammatory profiles. Metabolically healthy obesity (MHO) identifies individuals with obesity but a relatively preserved metabolic state. However, the criteria defining MHO remain inconsistent, and little is known about the gut microbiome (GM) features underlying this intermediate phenotype. Here, we aim to describe microbial structures contributing to metabolic health and disease. To do so, we analyzed the GM of 959 individuals classified as metabolically healthy non-obese (MHNO), MHO, metabolically unhealthy non-obese (MUNO), and metabolically unhealthy obese (MUO), using stool shotgun metagenomics. MHO subjects display intermediate anthropometric and biochemical profiles, with a GM composition and diversity in an in-between state among MHNO and MUO individuals. Network science analyses reveal that metabolic health, rather than obesity, drives microbial connectivity: MHNO and MHO individuals harbor more robust and functionally cohesive microbial networks, whose most influential nodes are focused toward SCFA production. In contrast, MUO and MUNO communities exhibit a dysbiotic state with reduced connectivity and increased influence of low-abundance, ectopic and potentially pro-inflammatory species resulting in a damaged, unstable microbial community network. These findings suggest that metabolic disorders disrupt microbial ecology beyond compositional shifts, emphasizing the need for systems-level approaches. Our findings show differences in microbial connectivity and association patterns across metabolic and obesity phenotypes, shedding light on how distinct microbial structures may contribute to metabolic health and disease.

## Introduction

Obesity is a chronic, multifactorial, noncommunicable disease defined by an excessive accumulation of fat^1,2^. World Health Organization (WHO) guidelines define obesity based on body mass index (BMI) alone, considering patients with BMI ≥ 30 kg/m^2^ as obese^3^. However, this metric does not accurately reflect adipose tissue distribution or muscle and fat mass proportions, which are key risk factors of cardiovascular disease. This has prompted the development of alternative frameworks to classify obesity patients, highlighting the relevance of the clinical rather than just the anthropometric component of obesity^2,4,5^. These underscore the nature of obesity as a heterogeneous condition, the limitations of BMI as a standalone measure of health, and the need to consider how excess adiposity affects health at an individual level.

Most of these initiatives view obesity as a progressive disease where health impairments can appear if excess weight is not managed correctly and even become disabling or life-threatening in further stages of the disease. In the initial stages of the disease, however, subjects would be asymptomatic, maintaining glucose homeostasis and adequate immune function, and without any functional impairments despite their increased body fat^1^. This phenotype has been termed metabolically healthy obesity (MHO). Currently, MHO is understood as an obesity phenotype where metabolic alterations caused by excessive adiposity are not present, and where insulin sensitivity and adipose tissue functionality are preserved^6^. MHO subjects are characterized by lower levels of ectopic fat storage, normal glucose metabolism, and lower inflammatory markers, all of which would provide them with protection against cardiovascular risk when compared to their metabolically unhealthy obese (MUO) counterparts^1,6^.

The exact markers that should be used to define MHO are still under debate. The BioShare-EU Healthy Obese Project, acknowledging the need for unified criteria, proposed a definition based on blood pressure and on serum concentrations of triglycerides, HDL cholesterol, blood glucose, and the absence of antihypertensive or glucose-lowering drug treatments^7^. Several longitudinal studies agree that it is a transient state that will eventually develop towards MUO if the excess weight is not treated, despite the variety in their definition of MHO^1,6,8^. Therefore, current research focuses on delaying this transition as much as possible to maintain patient quality of life, cardiometabolic health, and noncommunicable disease-free years^1,6^. This approach requires a thorough understanding of the mechanisms underlying MHO and its evolution towards MUO, as well as the markers that predict this transition.

Alterations in the gut microbiota (GM) have been associated with disruptions in immune regulation, systemic low-grade inflammation, and metabolic dysfunction, suggesting a potential mechanistic link between microbiota dynamics and the progression of metabolic impairments in obesity.^9^ Moreover, the GM has a bidirectional relationship with energy balance and weight management: it can causally influence both sides of the energy balance equation, while also being influenced by weight-modulating interventions such as diet, physical activity, surgery, and pharmacological approaches, as reviewed elsewhere^9^. Given its critical role in metabolic and inflammatory processes, understanding the interplay between GM composition and obesity phenotypes has become an area of active research.

Differences in the GM of obese subjects with different metabolic phenotypes have been explored in several studies^10–17^. These studies mostly view the MHO microbiome as an intermediate state between non-obese and MUO conditions: obesity would be the main driver of GM remodeling, with further changes occurring with the onset of metabolic disorders. However, differences in the definition of MHO and study design make it difficult to compare results and establish common signatures of the MHO microbiome. Many of these studies lack comparison with normal-weight^12,14,15,17^ or metabolically unhealthy subjects^16,17^, use experimental designs that do not address MHO specifically^10–12^, or focus on very specific populations, so that results might be difficult to generalize^13–15^. Nevertheless, they have yielded lists of potential microbial biomarkers for MUO or MHO by examining the direct associations between microbial features and the biochemical or anthropometric markers of obesity and metabolic health. This approach, however, overlooks the complex ecological networks and interactions within the GM, which play a crucial role in maintaining microbial community stability and host metabolism. By neglecting the broader ecological context, these studies may fail to capture microbial interactions and functional dynamics that contribute to metabolic health, potentially leading to an incomplete or misleading interpretation of the role the GM plays in obesity phenotypes.

Network science can be a useful tool to examine the ecological behaviors of microbial communities, providing results that are useful not only for biomarker identification but also from a community perspective. A wide variety of microbial ecosystems have been extensively studied through co-occurrence networks. Computationally inferred interactions have been experimentally validated in several settings, from the lung and skin myco- and microbiomes to a variety of environmental microbiomes, demonstrating how co-occurrence networks can be used to obtain biologically relevant information^18–21^. These methodologies can also be used to study ecosystem quality or the effect that different environmental conditions, including ecological stressors such as drought or climate warming, have on microbial communities^22–24^. In the human microbiome field, co-occurrence networks are being increasingly used to look for patterns in a variety of settings, describing microbial community features that may be associated with holt health or disease states ^18,25,26^. These studies show the promise that network-based methods hold for the study of human microbial communities.

Here, we propose to use network-based methods to provide an extensive characterization of GM structures, performed on the largest cohort of GM shotgun data focused on obesity and its related metabolic health states to date. Our dataset comprises 959 subjects classified as metabolically healthy obese (MHO), metabolically unhealthy obese (MUO), metabolically healthy non-obese (MHNO), and metabolically unhealthy non-obese (MUNO). We explore the GM not just through taxonomic profiling, based on microbial diversity and differential abundance analyses, but also by comparing its structural and functional organization among the four groups. By building and examining co-occurrence networks, we explored how microbial communities are organized and how their connectivity differs across metabolic phenotypes. Our findings show differences in microbial connectivity and association patterns across metabolic and obesity phenotypes, shedding light on how distinct microbial structures may contribute to metabolic health and disease.

## Materials and methods

### Data collection

#### Publicly available data

We searched the data available within the curatedMetagenomicData R package (version 3.8). This resource stores 93 publicly available human microbiome whole-genome shotgun datasets from different body sites, which have been processed with the same bioinformatics pipeline and whose metadata have been manually curated^27^. We searched for datasets with stool samples, with adult subjects that had not been treated with antibiotics recently, and where any of the following variables were available: body mass index, gender, blood glucose, triglycerides, HDL cholesterol, systolic or diastolic blood pressure, medication intake, or information regarding metabolic diseases. We further narrowed our search by discarding studies performed on Asian, African, or non-Westernized populations to avoid geographic variability, and by discarding subjects affected by conditions outside of our scope (e.g., colorectal cancer or celiac disease). The source publications of each dataset were accessed to look for inclusion criteria and for further variables that might not have been included in curatedMetagenomicData.

#### AI4Food project

We also included 98 samples from the AI4Food project^28,29^. This project was carried out on a cohort of 100 obese and overweight subjects (BMI ≥ 25 kg/m^2^) who went through a one-month-long weight loss intervention, during which lifestyle data was collected using diverse methodologies^28–30^. We classified subjects as MHNO, MHO, MUNO, or MUO at the beginning and final stages of the intervention and chose the healthiest stage for each subject. For instance, if a subject was MUO at baseline and their measurements after the intervention classified them as MHO, we used the stool sample and metadata collected after the intervention.

#### Phenotype assignment

Subjects were classified as metabolically healthy (MH) or unhealthy (MU) according to the BioSHaRE-EU Healthy Obese Project^7^. MH patients must comply with the following criteria: low fasting blood glucose (≤ 6.1 mmol/l or ≤ 100 mg/dl), low fasted serum triglycerides (≤ 1.7 mmol/l or ≤ 150 mg/dl), high HDL cholesterol concentrations (> 1.0 mmol/l or > 40 mg/dl in men and > 1.3 mmol/l or > 50 mg/dl in women), systolic blood pressure ≤ 130 mmHg and diastolic blood pressure ≤ 85 mmHg. Patients with T2D, impaired glucose transport, hypertension, hypercholesterolemia, or fatty liver disease, as well as those undergoing drug treatments against any of these disorders, were automatically considered MU regardless of their biochemical measurements. Further classification of the patients as MHNO, MHO, MUNO or MUO was performed based on body mass index following the WHO definition for obesity in Western adults (BMI ≥ 30 kg/m^2^)^3^.

### AI4Food data collection and preprocessing

#### Sample collection and DNA extraction

Stool samples from the AI4Food project were collected and frozen at −80 °C. DNA isolation was performed using the QIAamp Fast DNA Stool Mini Kit DNA extraction following the manufacturer’s instructions (QIAGEN, Hilden, Germany). Microbial analysis was performed by metagenomics shotgun sequencing on the NovaSeq 6000 Illumina platform (2 x 150 bp) with a coverage of approximately 6 GB per sample, equivalent to ∼40 million reads.

#### Taxonomic profiling

Metagenomic analyses for AI4Food samples were performed relying on the bioBakery computational environment and following the guidelines provided by curatedMetagenomicData to ensure dataset compatibility. Read trimming and removal of host reads were performed with the default configuration of the *read_qc* module from metaWRAP (version 1.2.1)^31^ using the hg38 version of the human genome for host read removal. The taxonomic profiling and quantification of relative abundances have been performed using MetaPhlAn4 (version 4.1)^32^ using the “mpa3” parameter and mapping against the mpa_v30_CHOCOPhlAn_201901 database to replicate curatedMetagenomicData’s pipeline.

### Batch effect correction

After merging the curatedMetagenomicData and AI4Food datasets, microbial species with an abundance above 0.01% in 10% of the samples were kept for further analyses. The MMUPHin R package (version 1.14)^33^ was used for batch effect correction caused by the study of origin while controlling for the effect of metabolic health and obesity. The effect of batch adjustment was evaluated based on the total variability in microbial profiles attributable to differences in the study of origin. This was done with a permutational multivariate analysis of variance (PERMANOVA) with 999 random permutations using the adonis2 function from the vegan R package (version 2.6-8)^34^.

### Microbial diversity analyses

Alpha diversity was estimated through the Chao1 index for richness and Shannon’s and (version 1.22)^35^. For beta diversity, the Aitchison distance, defined as the Euclidean distance between taxa after centered log-ratio data transformation, was calculated. Differences between groups were evaluated based on PERMANOVA with 999 permutations.

Differential abundance (DA) testing was performed with Analysis of Compositions of Microbiomes with Bias Correction 2 (ANCOM-BC2), an extension of the ANCOM-BC methodology that can be implemented in datasets with multiple groups (version 2.2.2)^36–38^. Multiple pairwise comparisons were performed while controlling for the mixed directional false discovery rate (mdFDR) using the Holm-Bonferroni procedure.

### Machine learning classifiers

The centered log-ratio (CLR) transformation was applied to batch effect-corrected relative abundances, which were used as input features in all machine learning models. Models were built using the caret R package (version 6.0-94)^39^ with curatedMetagenomicData abundances, using a 75/25% split for training and testing. Training was performed via 10-times 10-fold cross-validation while performing grid search for hyperparameter tuning based on Cohen’s Kappa score. Subsampling procedures based on upsampling or downsampling were tested but did not improve performance as measured by the area under the receiver operating characteristic curve (AUC).

#### Binary Models

Binary models were used to separate MH from MU subjects. Three algorithms common in the GM field^40^ were considered: Random Forest (RF), Support Vector Machines (SVM), and eXtreme Gradient Boosting (XGBoost). We used the ranger package (version 0.17)^41^ implementation of the RF algorithm. Hyperparameter tuning was used to choose the best values for the number of trees, the number of variables chosen at each split, and the minimal node size. The impurity decrease at each split was calculated via the Gini index criterion. Three SVMs with radial basis kernel functions were trained while performing hyperparameter tuning for the cost parameter *C* and sigma (kernlab package version 0.9-33)^42,43^. We also trained an XGBoost model using the xgboost R package (version 1.7.8.1)^44^. We first tuned learning rate, tree depth, and child node size, and proportions for row and column subsampling; and subsequently tuned regularization parameters alpha, lambda, and gamma. Cross-validation was performed for a maximum of 3000 iterations and with early stopping at 50 rounds. Logistic regression for classification was used as the objective function.

#### Multiclass Model

Since SVM was the best-performing algorithm in the binary case, we implemented an SVM model with a radial basis kernel function for multiclass classification, following the same strategy for model training and hyperparameter tuning as in binary models. To evaluate model performance, we calculated one-vs-rest receiver operating characteristic (ROC) curves, where each label (i.e., MHNO, MHO, MUNO and MUO) is compared against the other three.

### Co-occurrence networks

#### Network construction

Networks were generated using the Sparse InversE Covariance estimation for Ecological Association and Statistical Inference (SPIEC-EASI) method in “glasso” mode^45^. SPIEC-EASI builds networks in a compositionally aware manner, computing conditional independence between taxa instead of correlations. The SPIEC-EASI algorithm was run using the netConstruct function from the NetCoMi R package (version 1.1)^46^, using phyloseq (version 1.46)^47^ objects for each of the 4 groups as input. In the resulting network, nodes represent microbial species and edges represent species co-occurrence based on conditional independence. Keystone taxa were defined as nodes with degree and betweenness centrality greater than the 90 quartile using the netAnalyze function.

#### Network visualization

Networks were visualized in NetCoMi using the “spring” option, which uses the Fruchterman-Reingold algorithm to create a force-directed layout^48^. Force-directed layouts treat nodes in a network as if they were influenced by physical forces, which move the nodes into positions balancing attraction and repulsion. The objective is to achieve a layout where the distances between nodes are a good representation of edge weights between them.

#### Network structure

Network analyses were carried out in the NetworkX python library (version 3.4.2)^49^. The largest connected component (LCC) of each network was used for subsequent analyses. The SPIEC-EASI algorithm can generate more than one subnetwork, each of which is called a connected component. The LCC is the connected component containing the most nodes. The topological parameters that we have analyzed are: network order and size, edge density, number of connected components, node degree, average shortest path length, node betweenness centrality, and closeness centrality. Network order and size reflect the number of nodes and edges respectively. Edge density is the ratio of the number of edges in a network versus the maximum number of possible edges it can contain. Node degree represents the number of connections that each node shares with the rest of the network. We also performed *k*-core decomposition analysis. A *k*-core is a subgraph (i.e., a set of nodes within the network) where all nodes share at least *k* edges between them.

#### Distances in weighted co-occurrence networks

Node betweenness centrality, average shortest path length, and closeness centrality are measurements obtained from the shortest paths in the network. The average shortest path length is the mean shortest distance between all pairs of nodes in a network. Node betweenness centrality measures the proportion of shortest paths within the network that go through a specific node. Node closeness centrality measures how reachable other nodes in the network are, and is calculated as the reciprocal of the sum of the shortest path lengths to all other nodes in the graph. The shortest path between two nodes is the route with the fewest edges between them. In weighted networks, shortest path lengths are calculated considering these weights, and the shortest path might not be the one with the least edges, but the less costly one. NetworkX interprets weights as distances: edges with higher weights will have a higher associated cost. However, in conditional independence networks, higher weights should result in lower costs, since they represent closer relations between taxa. We have adjusted our data by subtracting each conditional independence value from 1. This transformation ensures that higher values (i.e., longer distances) are assigned to lower correlations, aligning with NetworkX’s underlying assumptions.

#### Network comparisons

We performed Kruskal-Wallis rank sum tests to analyze local properties, i.e., where values can be calculated for each node: node degree, betweenness centrality, and closeness centrality. We also performed this test on shortest path lengths between all pairs of nodes. The remaining measurements refer to global network properties, yielding a single value per network.

#### Network robustness and stability

To analyze network robustness, we implemented a framework to further analyze network stability based on targeted and random attacks. First, negative edges were removed from the network to only consider positive microbe-microbe associations. Network nodes were sorted according to the property of interest (i.e., degree, betweenness centrality, decreasing mean abundance, and increasing mean abundance) and the node with the maximum value for said property was removed from the network. Then, the number of nodes of the remaining LCC was measured. This process was repeated until all nodes were disconnected from each other. In the case of random attacks, nodes were chosen randomly using the choice method from the random module, and the process was repeated 1000 times for reproducibility. To measure resistance against attacks, we coined a measure named node removal 50 (NR_50_), representing the percentage of nodes that need to be removed from the network so that the resulting LCC has half its original nodes.

### Statistical analyses

All statistical analyses were performed in R version 4.3.2. Quantitative variables were compared between the 4 groups using the Kruskal-Wallis rank sum test, and post-hoc comparisons were made via two-sided Dunn’s test. Benjamini-Hochberg’s false discovery rate (FDR) was used to correct for multiple comparisons.

### Data and code availability

All code used for microbial profiling, network construction and analysis, and machine learning is available in a GitHub repository (https://github.com/blacruz17/MHOmicrobiome). AI4Food microbial samples are available at the European Nucleotide Archive with accession code PRJEB87701. AI4Food patient metadata are provided in Supplementary Table 1. Microbial abundances and sample metadata for the remaining studies can be accessed via curatedMetagenomicData or their source publications, cited in the main text. Supplementary Table 2 provides the subject IDs and phenotype classifications for each of these studies.

## Results

### Selection and curation of microbiome datasets for metabolic health and obesity analyses

We mined the 93 human microbiome datasets available in the R package curatedMetagenomicData for studies where subjects could be classified according to their metabolic health and obesity status. To do so, we searched for studies with stool samples from adult subjects not taking antibiotics, and where at least one of our variables of interest, described in the Methods section, was available (Supplementary Figure 1). After this search, we were able to keep 4 studies where our eligibility criteria were met and with enough information to classify subjects as MHNO, MHO, MUNO or MUO, either on the curatedMetagenomicData package or on the source publication: MetaCardis_2020_a^50^ (n = 629 subjects), KarlssonFH_2013^51^ (n = 145), FengQ_2015^52^ (n = 61) and HMP_2019_t2d^53^ (n = 27). We also included 98 samples from the AI4Food study^28,29^. The number of subjects classified in each phenotype per study is shown in Table 1.

**Table 1.**
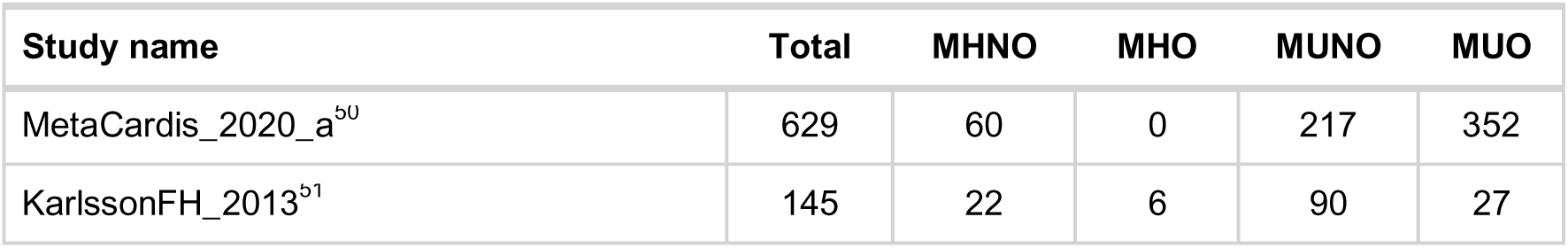
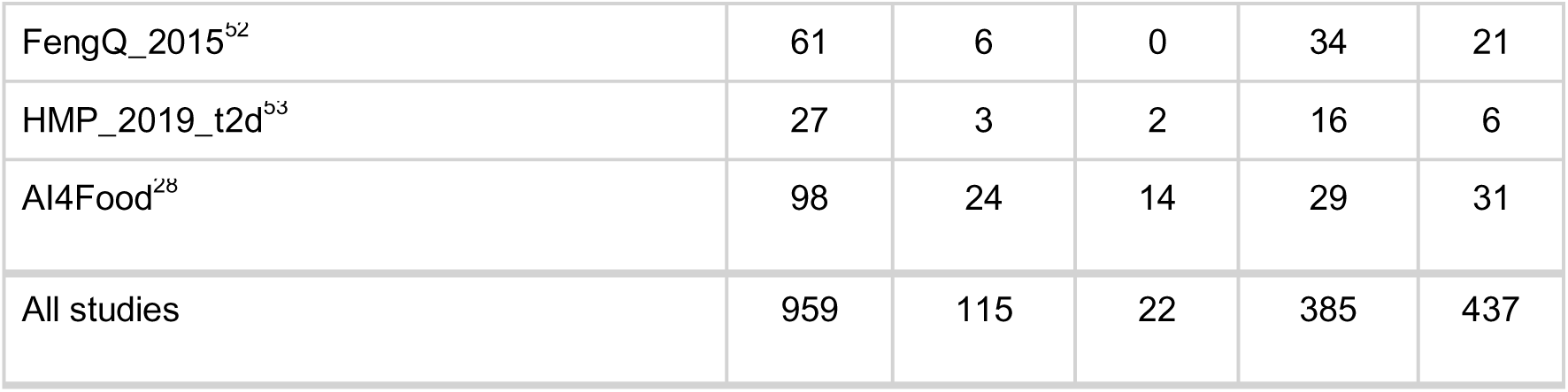
Datasets used: number of samples per study and group.

### MHO subjects have lower adiposity and distinct metabolic-inflammatory profiles *vs.* MUO and non-obese groups

First, we analyzed subjects’ metadata to evaluate their metabolic and inflammatory profiles, shown in Table 2 and Supplementary Figure 2. MUNO subjects are the oldest, with a mean age of 67, while the remaining groups have mean ages of 54 (MHNO, p < 0.001), 58 (MHO, p < 0.01) and 61 (MUO, p < 0.001). This agrees with current research viewing age as an accelerator in the MHO-to-MUO transition.^1^ MHNO subjects, as expected, have the lowest BMI among the four groups (23 kg/m^2^, p < 0.001), while the MUNO group has a mean BMI classified as overweight (27 kg/m^2^). Moreover, 132 MUO subjects (30.1%) reach a BMI ≥ 40 kg/m^2^, while no MHO subjects reach this threshold, which is considered grade III or high-risk obesity by WHO^1^ and enough to consider excessive fat accumulation by more recent guidelines^2,5^. Relevant differences were found as well in other anthropometric measurements used to evaluate adiposity, such as waist circumference or waist/hip ratio (WHR)^2,5^, which are higher in MUO (waist circumference: 106 cm, WHR: 0.92) than in MHO subjects (waist circumference: 100 cm, p < 0.05; WHR: 0.86, p < 0.001). This aligns with the view of obesity as a continuum, where MHO subjects show lower BMI and adiposity levels and MUO patients have higher levels of excess fat, reflected by the presence of high-risk obesity and larger waist circumference and WHR values.

**Table 2.** Population characteristics.

| Characteristic | N | MHNO<br>N = 115<sup>1</sup> | MHO<br>N = 22<sup>1</sup> | MUNO<br>N = 385<sup>1</sup> | MUO<br>N = 437<sup>1</sup> | p-value<sup>2</sup> | adjusted<br>p-value<sup>3</sup> |
| --- | --- | --- | --- | --- | --- | --- | --- |
| Age (years) | 606 | 54 (42, 66) | 58 (38, 70) | 67 (60, 70) | 61 (48, 69) | <0.001 | <0.001 |
| Sex (% female) | 959 | 79 (69%) | 20 (91%) | 194 (50%) | 266 (61%) | <0.001 | <0.001 |
| Medication intake<br>(number of patients)* | 872 | 0 (0%) | 0 (0%) | 244 (73%) | 292 (71%) | <0.001 | <0.001 |
| BMI (kg/m<sup>2</sup>) * | 959 | 23 (22, 26) | 33 (31, 35) | 27 (25, 29) | 36 (32, 42) | <0.001 | <0.001 |
| Hip circumference<br>(cm) | 141 | 107 (103, 111) | 118 (115, 120) | 105 (101, 109) | 114 (111, 120) | <0.001 | <0.001 |
| Waist circumference<br>(cm) | 290 | 86 (80, 92) | 100 (93, 104) | 91 (84, 97) | 106 (100, 113) | <0.001 | <0.001 |
| Waist/hip ratio | 286 | 0.85 (0.80, 0.88) | 0.86 (0.81, 0.87) | 0.87 (0.82, 0.93) | 0.92 (0.88, 0.97) | <0.001 | <0.001 |
| Systolic blood pressure<br>(mmHg)* | 97 | 73 (69, 78) | 76 (72, 79) | 75 (71, 80) | 85 (81, 89) | <0.001 | <0.001 |
| Diastolic blood<br>pressure (mmHg)* | 97 | 111 (104, 118) | 119 (114, 126) | 119 (107, 131) | 137 (125, 144) | <0.001 | <0.001 |
| Glucose (mg/dL) * | 323 | 87 (81, 95) | 82 (78, 90) | 102 (88, 115) | 101 (89, 123) | <0.001 | <0.001 |
| HbA1c (%) <sup>4</sup> | 549 | 5.40 (5.20, 5.50) | 5.52 (5.35, 5.80) | 5.70 (5.44, 6.00) | 5.70 (5.40, 6.00) | <0.001 | <0.001 |
| HOMA-IR<sup>5</sup> | 128 | 1.31 (0.92, 1.58) | 2.12 (1.09, 2.90) | 1.74 (1.15, 2.49) | 3.02 (1.85, 4.35) | <0.001 | <0.001 |
| Insulin (μU/mL) | 273 | 6 (5, 7) | 9 (6, 14) | 8 (6, 12) | 14 (9, 21) | <0.001 | <0.001 |
| Triglycerides (mg/dL) * | 572 | 75 (57, 92) | 81 (61, 101) | 94 (74, 143) | 121 (93, 159) | <0.001 | <0.001 |
| Cholesterol (mg/dL) | 262 | 215 (187, 235) | 201 (186, 217) | 208 (181, 225) | 199 (175, 236) | 0.5 | 0.5 |
| HDL (mg/dL) * | 320 | 70 (58, 85) | 62 (53, 70) | 61 (49, 77) | 53 (44, 65) | <0.001 | **<0.001** |
| LDL (mg/dL) <sup>6</sup> | 571 | 126 (104, 144) | 123 (105, 137) | 122 (99, 148) | 125 (103, 151) | 0.8 | 0.8 |
| TNF-α (U/mL) <sup>7</sup> | 98 | 3.13 (2.40, 5.19) | 5.18 (3.93, 6.79) | 5.58 (4.46, 6.50) | 4.88 (3.42, 6.37) | 0.006 | **0.009** |
| TNF-α (ng/L) <sup>7</sup> | 145 | 1.82 (1.33, 2.31) | 1.71 (0.94, 2.90) | 1.85 (1.56, 2.47) | 2.18 (1.88, 2.94) | 0.2 | 0.2 |
| C-reactive protein (mg/dL) | 158 | 0.10 (0.03, 0.30) | 0.16 (0.03, 0.41) | 0.20 (0.10, 0.36) | 0.20 (0.10, 0.44) | 0.024 | **0.030** |
| Adiponectin (μg/dL) | 242 | 14 (9, 17) | 13 (9, 15) | 12 (9, 16) | 10 (8, 14) | 0.038 | **0.047** |
| Albumin (g/dL) | 118 | 4.50 (4.30, 4.60) | 4.35 (4.20, 4.60) | 4.35 (4.20, 4.60) | 4.40 (4.40, 4.60) | 0.084 | 0.10 |

**MHNO:** metabolically healthy non-obese. **MHO:** metabolically healthy obese. **MUNO:** metabolically unhealthy non-obese. **MUO:** metabolically unhealthy obese. **1** n (%); Median (Q1, Q3) **2** Pearson's Chi-squared test; Kruskal-Wallis rank sum test **3** False discovery rate correction for multiple testing **4** Hemoglobin A1C **5** Homeostatic Model Assessment for Insulin Resistance **6** Low-density lipoprotein **7** Tumor necrosis factor alpha. \*: variables used for phenotype assignment.

The four groups show differences in all markers used for metabolic health stratification. When compared to the MUO group, MHO subjects show lower triglycerides (81 *vs.* 121 mg/dL, p < 0.001), glucose (82 *vs.* 101 mg/dL, p < 0.001), and blood pressure (BP) (systolic BP: 76 *vs* 85 mmHg, p < 0.05; diastolic BP: 119 *vs.* 137 mmHg, p < 0.01), reflecting a metabolic state relatively unaffected by excess fat. This is accompanied by lower values in the glycemic profile, reflected in the HOMA index (2.12 *vs* 3.02, p < 0.05) and insulin concentrations (9 *vs.* 14 µU/mL, p < 0.05). However, this phenotype is not completely benign, as shown by the heightened glycemic parameters relative to the MHNO group (HbA1c: 5.52 *vs.* 5.40 %, p < 0.05; insulin: 9 vs. 6 µU/mL, p < 0.01; HOMA: 2.12 *vs.* 1.31, p < 0.05). This highlights MHO as an intermediate condition, where metabolic parameters related to glucose metabolism show deviations compared to normal-weight subjects. These parameters worsen when this condition evolves towards MUO, with a decline in glycemic and lipidic biochemical profiles.

Finally, differences in inflammatory profiles were also found: TNF-α levels (U/mL) are the lowest in the MHNO group (3.13 U/mL, p < 0.01); while C-reactive protein (CRP) levels are higher in MUNO and MUO (0.20 mg/dL in both cases), than in the MHNO group (0.10 mg/dL, p < 0.05), which do not show differences with MHO subjects (0.16 mg/dL). This aligns with a view of MHO as an intermediate profile between MHNO and MUO, where we can observe inflammatory patterns related to obesity, reflected by an increase in TNF-α, that are accompanied by an increase in CRP in subjects with more extreme metabolic phenotypes (MUNO, MUO). In summary, MHO subjects show a worsened metabolic-inflammatory condition when compared to their normal-weight counterparts, but not as severe as in MUO individuals.

### Metabolically unhealthy gut microbiomes are less diverse and richer in pro-inflammatory microbes compared with MHNO communities

Alpha diversity was evaluated through the Chao1 index for species richness and Simpson’s and Shannon’s indices for diversity (Figure 1A). MUO subjects had the lowest alpha diversities, showing significant differences with their non-obese counterparts (MUNO) in all three indices (p < 0.001, p < 0.05, and p < 0.001 for Chao1, Simpson’s, and Shannon’s indices respectively). We also found significantly lower indices when compared to MHNO subjects (p < 0.001, p < 0.01, and p < 0.001 for Chao1, Simpson’s, and Shannon’s indices respectively). Then, we calculated the Aitchison distance matrix as a measure of beta diversity, looking for differences in community structure and composition between the four groups. Even though the 4 populations were different (*p* < 0.001, PERMANOVA), they could not be separated visually in the ordination plots (PCoA, Fig. 1B).

**Figure 1.**
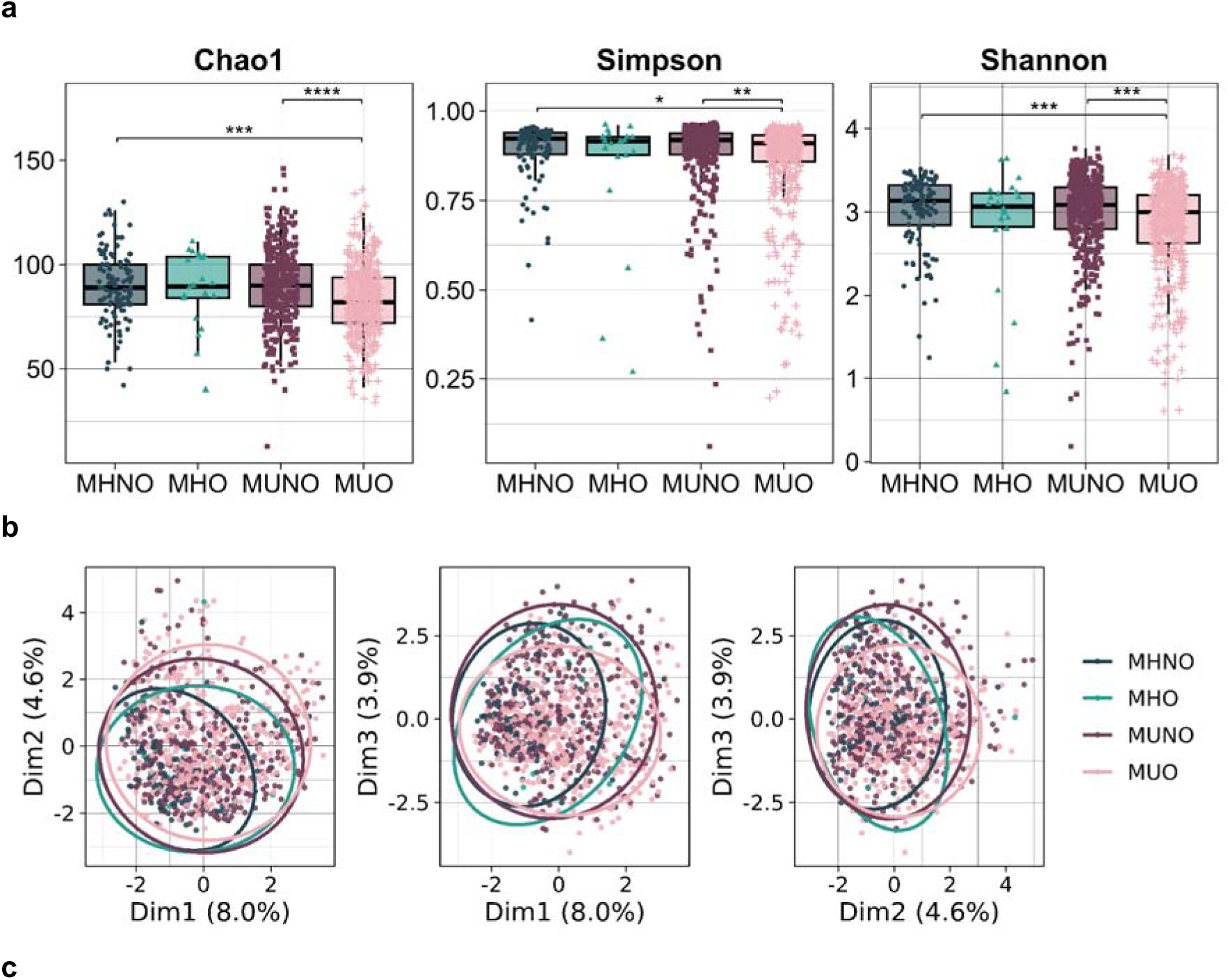
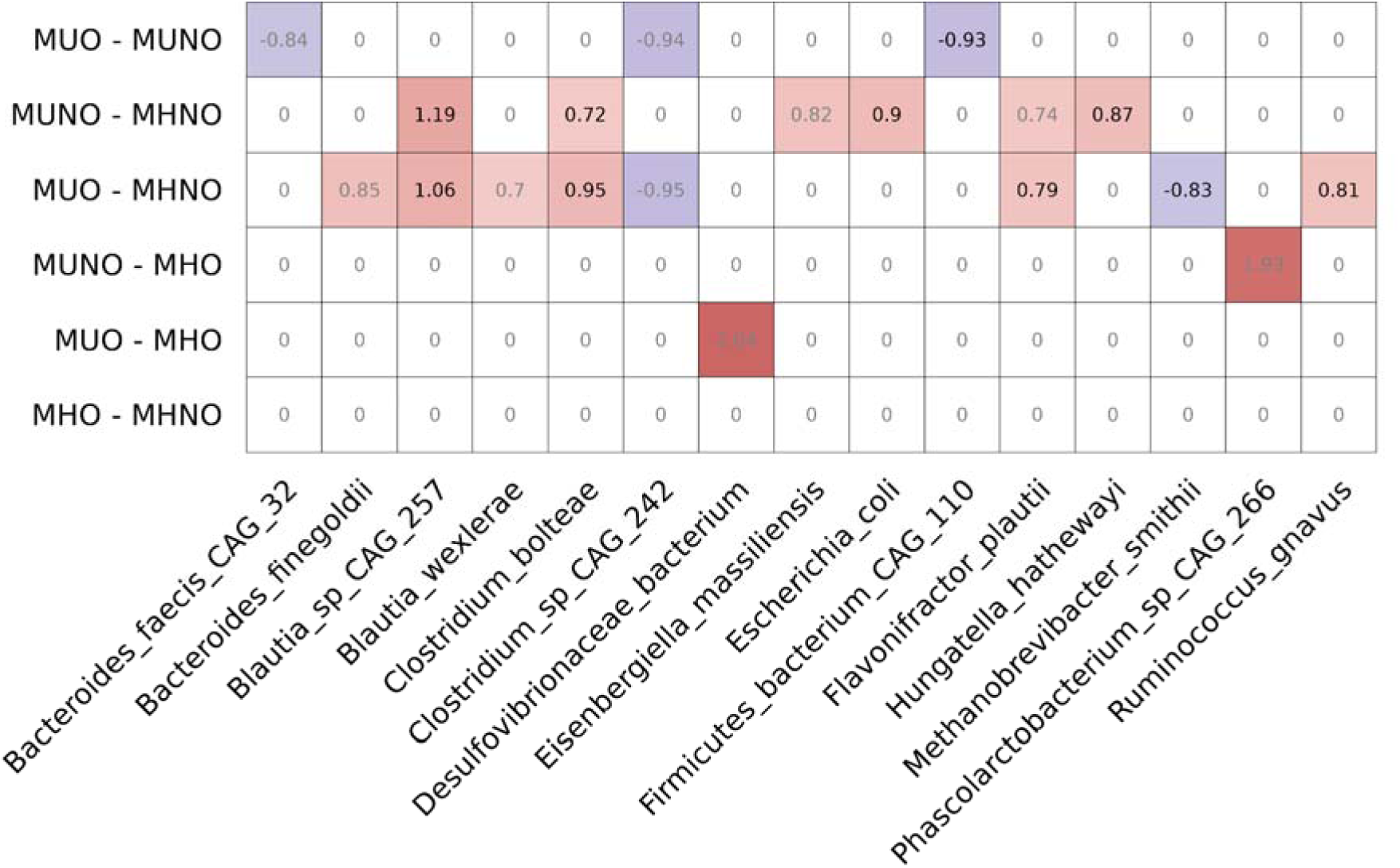
Gut microbiome exploration. **a)** Alpha diversity analysis. The Chao1 index is shown for species richness. Shannon’s and Simpson’s indices were chosen for species diversity. The Kruskal-Wallis rank sum test was used to evaluate differences (p values: Chao1 < 0.001, Simpson = 0.004, Shannon < 0.001). Pairwise comparisons were tested with Dunn’s post-hoc test. All p-values were FDR-corrected. **b)** Beta diversity analysis. Aitchison’s distances between samples were calculated and PCoA was chosen for graphical representation. Differences between groups were tested with PERMANOVA (999 permutations). Ellipses represent 95% confidence intervals. **c)** Differential abundance analysis. Analysis of Compositions of Microbiomes with Bias Correction (ANCOM-BC2) was used to determine differentially abundant taxa in multiple pairwise comparisons. Cells with values different from 0 represent comparisons with significant differences after false discovery rate (mdFDR) correction. Values and colors in the heatmap cells represent log fold-change (LFC) values.

We then looked for microbial species that could serve as biomarkers for each condition by performing differential abundance (DA) testing between groups. This returned a total of 15 taxa with significant differences (Figure 1C), many of which showed differential abundances when comparing the MHNO community to the MUO or MUNO groups. At the functional level, markers of metabolically unhealthy groups have been associated with increases in the pro-inflammatory metabolite trimethylamine N-oxide (*Bacteroides finegoldii*^54^); immune modulation (*Clostridium bolteae*^55^*, Flavonifractor plautii*^56^); and with the onset of colorectal cancer (CRC) (*F. plautii*^57–59^*, Hungatella hathewayi*^60,61^) or type 2 diabetes (*Clostridium bolteae*^62^). The increase in *Firmicutes bacterium CAG 110* in MUNO compared to MUO patients might be related to its ability to synthesize beneficial unsaturated fatty acids^63,64^. Finally, MHNO subjects show higher levels of the longevity marker *Methanobrevibacter smithii*^65^. These results suggest that the transition from MHNO to MUO, with MHO and MUNO as intermediate stages, involves changes in GM composition, characterized by the loss of microbial diversity and by the proliferation of microbes contributing to inflammation and metabolic dysregulation.

### Machine learning models could not discriminate metabolically healthy and unhealthy subjects

Differential abundance methods identify microbes whose abundances change between phenotypes but fail at identifying more complex patterns, which could serve as microbial signatures for metabolic disorders in normal weight or obesity settings. To study these complex patterns, we built several machine learning classifiers to solve two different classification problems: first, the comparison of metabolically healthy (MH) and metabolically unhealthy (MU) subjects; second, a multiclass problem with MHNO, MHO, MUNO, and MUO labels. We trained binary models based on Random Forest (RF), Support Vector Machines (SVMs), and eXtreme Gradient Boosting (XGBoost). The three models had a modest area under the receiver operating characteristic (ROC) curve (AUC), all between 0.60 and 0.70 (Figure 2A). For multiclass classification, we trained an SVM model, since it showed the highest AUC in the previous experiment (0.68). To evaluate its performance, we calculated

**Figure 2.**
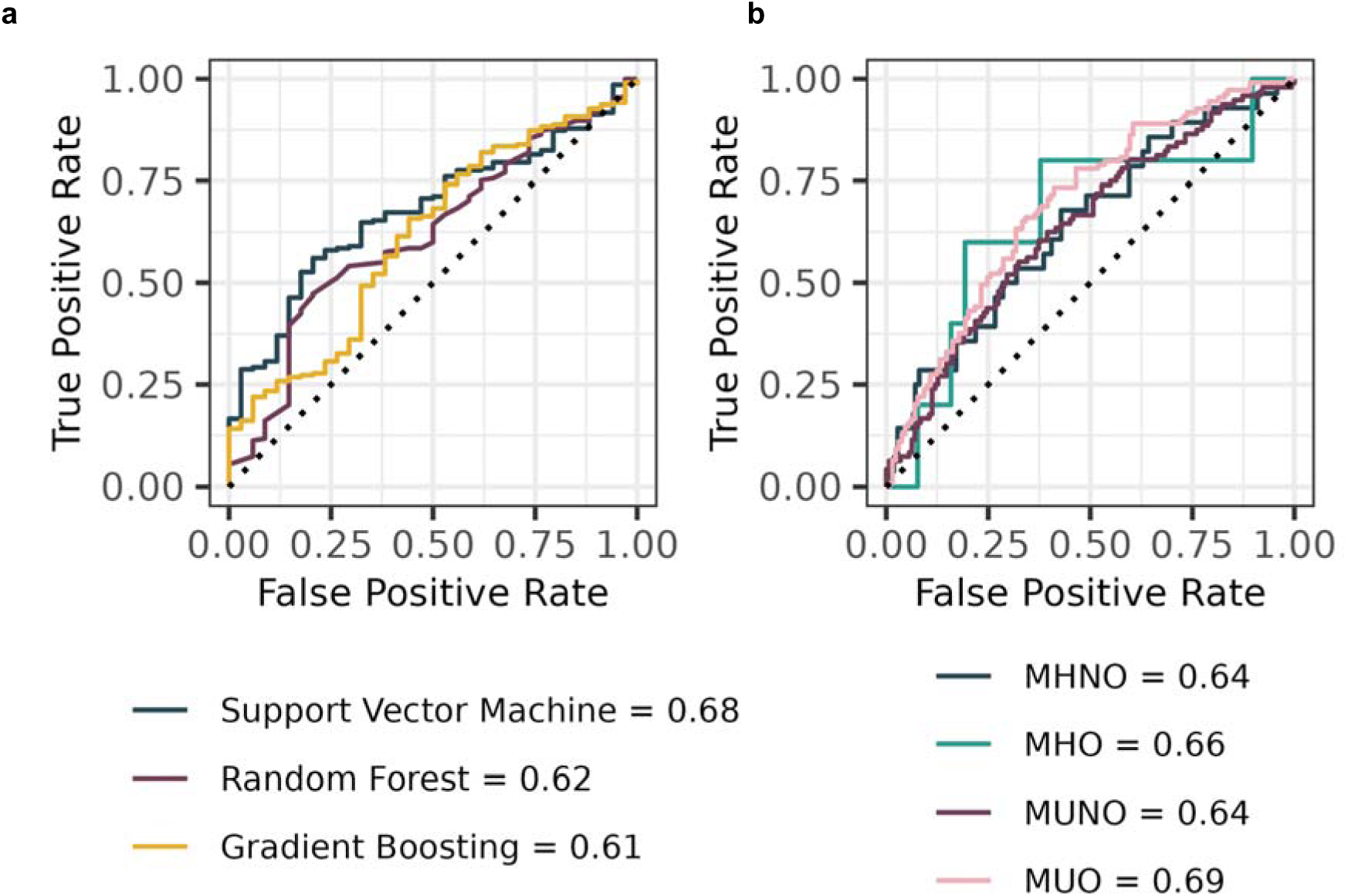
Machine learning models. Receiver operating characteristic (ROC) curves for the binary and multiclass models. Legends show area under the curve (AUC) values for the test set. **a)** ROC curves for binary models including Support Vector Machine, Random Forest and eXtreme Gradient Boosting. **b)** Multiclass model results for a SVM using a radial basis kernel function. One-vs-all ROC curves, as well as their AUCs, are shown.

One-*vs*-Rest ROC curves, where each class is compared against all others. The MUO class shows the highest AUC (0.69), while non-obese subjects are the most difficult to separate from the others (AUCs of 0.64 in both MHNO and MUNO cases). However, the AUCs are still low, with none of them reaching 0.70.

Our results indicate that the use of abundance-based data for patient classification yields limited predictive power. This can be attributed to the low number of subjects in healthy metabolic phenotypes (MHNO, MHO) and to the class imbalances among groups, which could not be compensated by subsampling approaches. These constraints underscore the need to address our problem from an ecological perspective, shifting the focus from differences in individual microbes to the broader structure and dynamics of the gut microbiota. To address this, we explored network-based approaches, which go beyond identifying differentially abundant microbial taxa or complex machine learning-derived patterns. Instead, these methods allow us to capture the ecological interactions within the microbiota, providing a comprehensive view of microbial community structure and how microbial interactions may contribute to metabolic and obesity phenotypes.

### Gut microbiota structure is reshaped in the transition from metabolically healthy normal-weight to metabolically unhealthy obesity

Microbial co-occurrence networks are widely used to represent different microbial communities and to better understand how these communities are structured. In these graphs, nodes represent microbial species and edges depict inferred interactions between them. To explore how obesity or metabolic disease shape the GMs of MHNO, MHO, MUNO, and MUO subjects, we built co-occurrence networks (CNs) for each phenotype.

The four resulting CNs are composed of 340 (MHNO), 289 (MHO), 334 (MUNO) and 326 (MUO) nodes. Network visualizations can be seen in Figure 3 (panels A-D), while network and node topological properties are shown in Table 3. Networks from MHNO (Fig. 3A) and MHO (Fig. 3B) subjects consist of a single connected component, where all microbes are connected to the rest of the community. However, in the case of MUNO (Fig. 3C) and MUO (Fig. 3D) communities, even though most microorganisms are part of the networks’ largest connected component (LCC), there are a series of nodes that are isolated from it, representing 6.48% and 9.89% of nodes respectively. Moreover, these graphs have lower 10^-2^, MHO: 1.92 x 10^-2^), indicating a higher number of connections in metabolically healthy microbiomes. This is accompanied by higher degrees in the MHNO (mean degree: 5.74) and MHO (5.54) CNs than in MUNO (4.57) and MUO (4.26) graphs (p < 0.001). This first glance into the different CNs already indicates differences in microbial interactions among the four phenotypes, where microbes in metabolically diseased communities are less connected to each other and form smaller communities than in metabolically healthy GMs.

**Figure 3.**
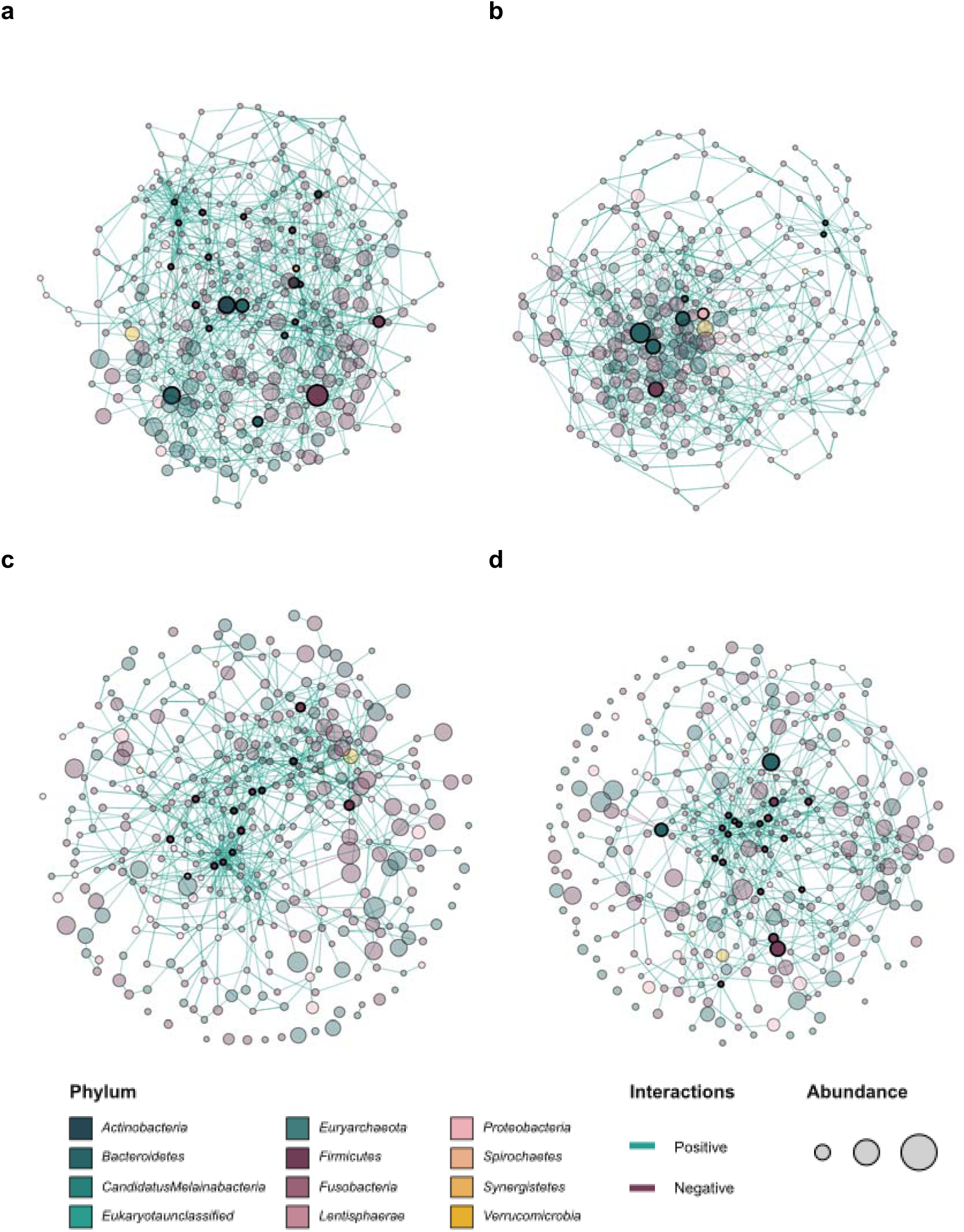
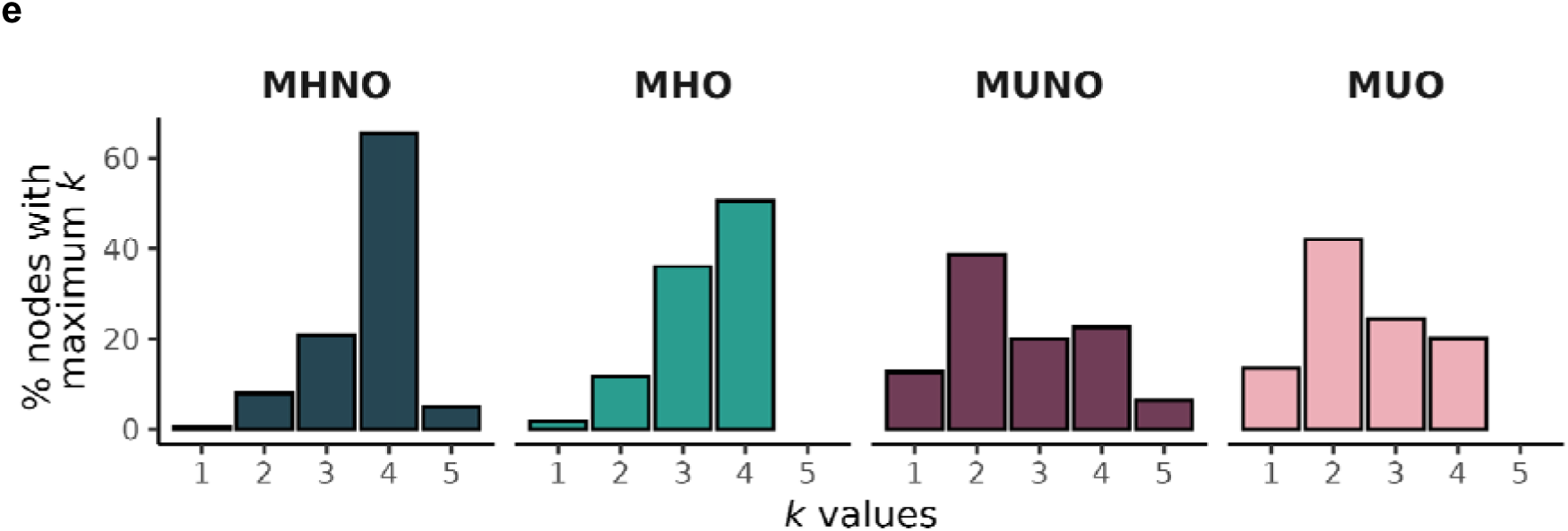
Co-occurrence networks. Networks obtained from the SPIEC-EASI algorithm in MHNO (**a**), MHO (**b**), MUNO (**c**), and MUO (**d**) groups. Node position was calculated using a force-directed layout based on the Fruchterman-Reingold algorithm. Nodes are colored based on their phyla. Edge colors represent conditional independence signs. Node size is scaled according to normalized relative abundances. Highlighted nodes represent keystone taxa. **e)** *K*-core distribution. Values are shown as the percentage of total nodes with maximum *k*-cores at different values of *k*.

**Table 3.**
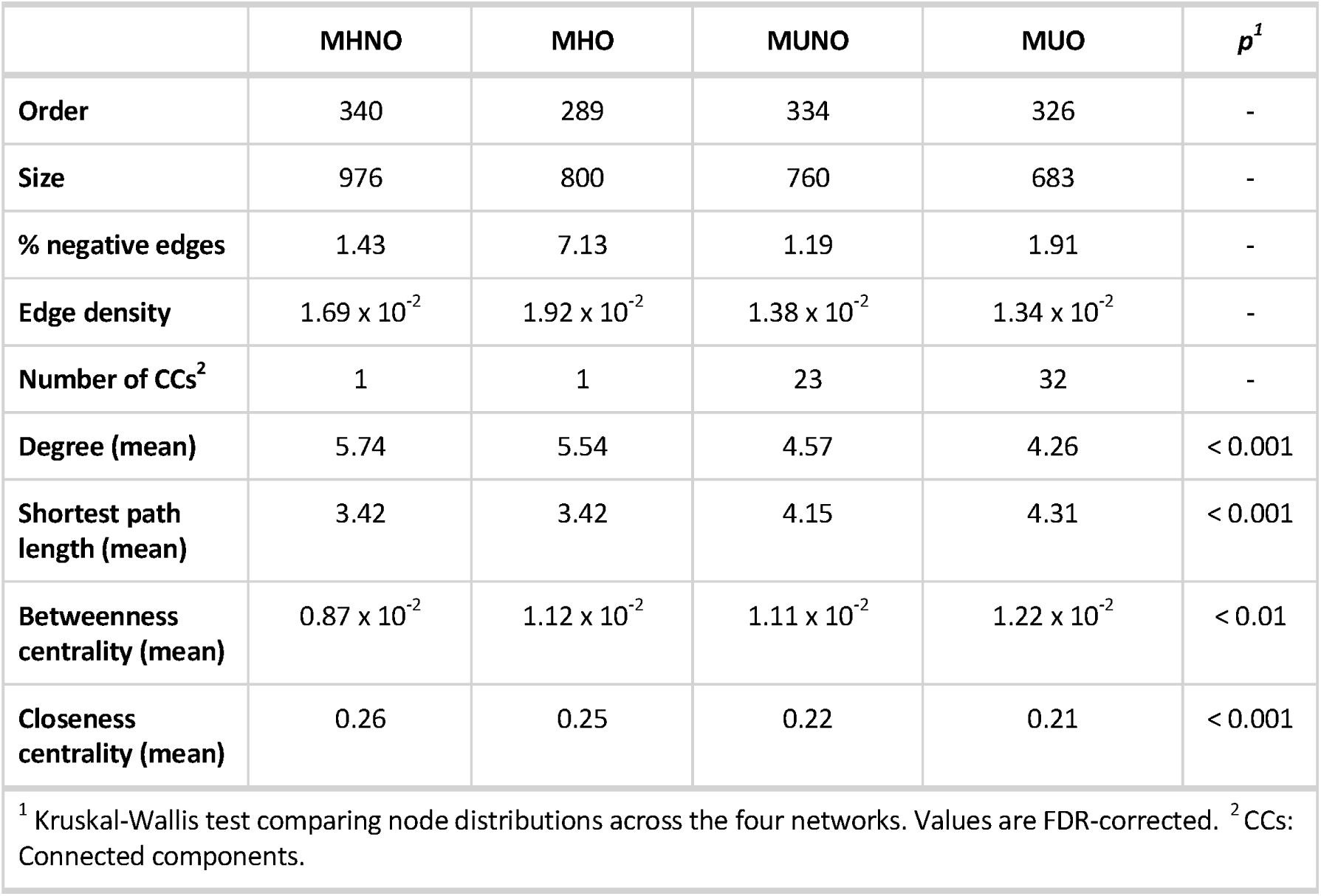
Summary of network topological properties. In the MUNO and MUO cases, all properties except order and size are given for the largest connected component.

We also calculated average shortest path lengths, which are shorter in MHNO and MHO communities (3.42 in both cases). MUNO subjects show an intermediate value for this metric (4.15), with the MUO graph showing the highest average shortest path length (4.31) (p < 0.001). In line with this, the MHNO CN has the lowest mean betweenness centrality (0.87 x 10^-2^) and the largest mean closeness centrality (0.26), the MUO CN shows an opposite trend (mean betweenness centrality: 1.22 x 10^-2^, mean closeness centrality: 0.21), and MHO and MUNO communities have intermediate values. These values suggest that information is distributed more evenly in the MHNO CN, which relies less heavily on central nodes and has shorter distances that are easier to travel, than in the MUO community, with MHO and MUNO CNs falling in an intermediate range. This might indicate that metabolic disease and obesity disrupt microbial community organization, reducing the efficiency of communication among its members. As the microbial network is reshaped and becomes increasingly dependent on central nodes, signaling and metabolite transmission would become less efficient, potentially contributing to metabolic dysfunctions observed in these conditions.

To further explore how connections are distributed, and how networks rely on central nodes, we visualized *k*-core distributions in each network (Figure 3E). Networks where most nodes have higher *k* values are more robust, which can be indicative of higher biological stability due to better cooperation among members of the community. In metabolically healthy networks, *k*-core distributions are shifted towards higher values, especially in the MHNO case, where 66% of nodes belong to 4-cores (MHNO) (Figure 3E). This percentage falls to 50% in the MHO case and is the lowest in MUNO and MUO communities, where 23% and 20% of nodes belong to 4-cores respectively. In these two CNs, *k*-core distributions are more evenly spread and tend towards lower values, especially in the MUO case. This indicates that metabolically healthy networks are more tightly connected than metabolically diseased microbial communities, aligning with the network topology results. In metabolically healthy contexts, microbes would form highly connected functional guilds (*k-*cores), further facilitating their cooperation.

### Keystone taxa composition shifts from short-chain fatty acid producers to ectopic microbes in metabolically diseased communities

Nodes with high degree, also called hubs, or betweenness, called bottlenecks, reflect highly influential nodes within a network. In microbial co-occurrence networks, these two properties are commonly used to define “keystone taxa”^66^: microbial species that are essential to maintain community structure and function, regardless of their abundance. Keystone taxa for our CNs are shown in Table 4.

**Table 4.**
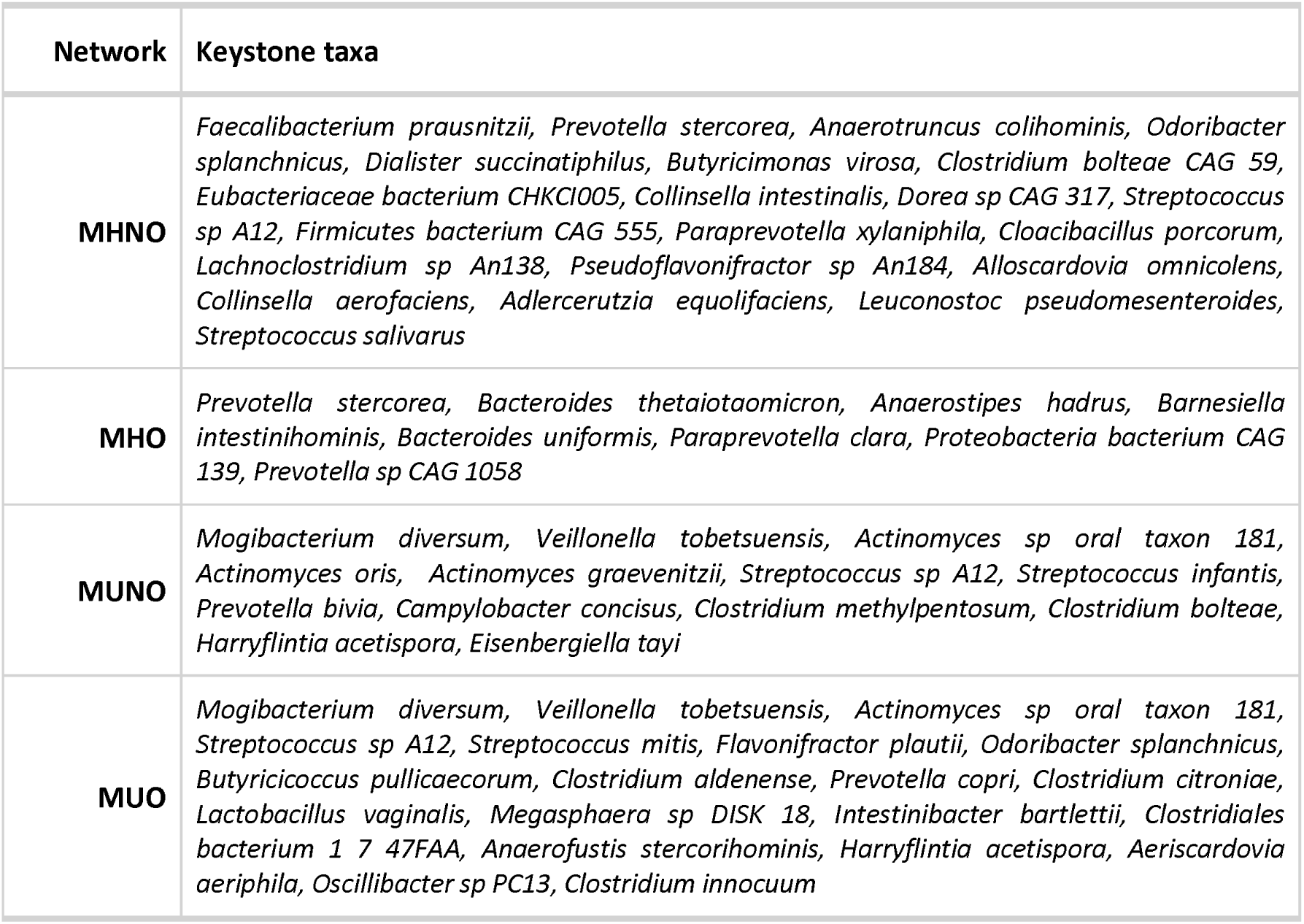
Keystone taxa from each microbial community.

Among the MHNO and MHO keystone taxa, we found prevalent members of the human GM (*Faecalibacterium prausnitzii*^67^*, Prevotella stercorea*^68^*, Bacteroides thetaiotaomicron*^69^) together with several short-chain fatty acid (SCFA) producers (*F. prausnitzii*^67^*, Anaerotruncus colihominis*^70^*, Odoribacter splanchnicus*^71^*, Dialister succinatiphilus*^72^*, Butyricimonas virosa*^73^*, Anaerostipes hadrus*^70,74^ and *Barnesiella intestinihominis*^72^). SCFAs produced in the gut regulate host metabolic and immune processes and thus might prevent metabolic dysregulation and low-grade inflammation occurring in obesity and metabolic disorders^9^. Other microbes with anti-inflammatory properties, known for their beneficial effects against irritable bowel disease (*B. thetaiotaomicron*^75^, *Bacteroides uniformis*^76^), were also detected.

Remarkably, we identified several members of the oral microbiome (*Mogibacterium diversum*^77^*, Veillonella tobetsuensis*^78^*, Campylobacter concisus*^79^*, Actinomyces spp.*^80,81^, and *Streptococcus spp.*^81,82^) among MUNO and MUO keystone taxa. Microbial translocation from the mouth to the gut across the oral-gut axis, which does not happen in healthy patients, has been described in inflammatory contexts, including IBD and CRC^83–85^. Translocation of these microbes, which would form consortia with colonic aerobic bacteria^85^, can have adverse effects on host health, including gut barrier disruption and expression of virulence factors^83,84^. *F. plautii* is also involved in CRC onset^57–59^, as well as in immune modulation^56^. SCFA producers (*O. splanchicus*^71^) and candidate probiotics (*Butyricicoccus pullicaecorum*^86^), necessary to maintain GM function even in dysbiotic communities, are also present.

Changes in microbial communities related to metabolic disorders involve a shift in keystone taxa that sustain community structure: while MHNO and MHO communities are dominated by SCFA producers, the GM of MUNO and MUO subjects is characterized by the presence of ectopic microbes from the oral microbiota, which may negatively impact host health.

### Metabolically healthy microbial communities form more robust structures than metabolically diseased phenotypes

After defining CN properties and keystone taxa, we aimed to compare community stability based on how graphs behave if their nodes are removed: we hypothesize that less stable networks will decay faster and lose their structure; while resilient microbial communities will not be as affected to such an extent and would require more attacks to be destabilized. For a quantitative measure of network stability, we devised node removal 50 or NR_50_, a measure reflecting the percentage of nodes that need to be removed from a network to reduce its number of nodes by half. Our methodological framework, detailed in the Methods section, is summarized in Figure 5A.

**Figure 5.**
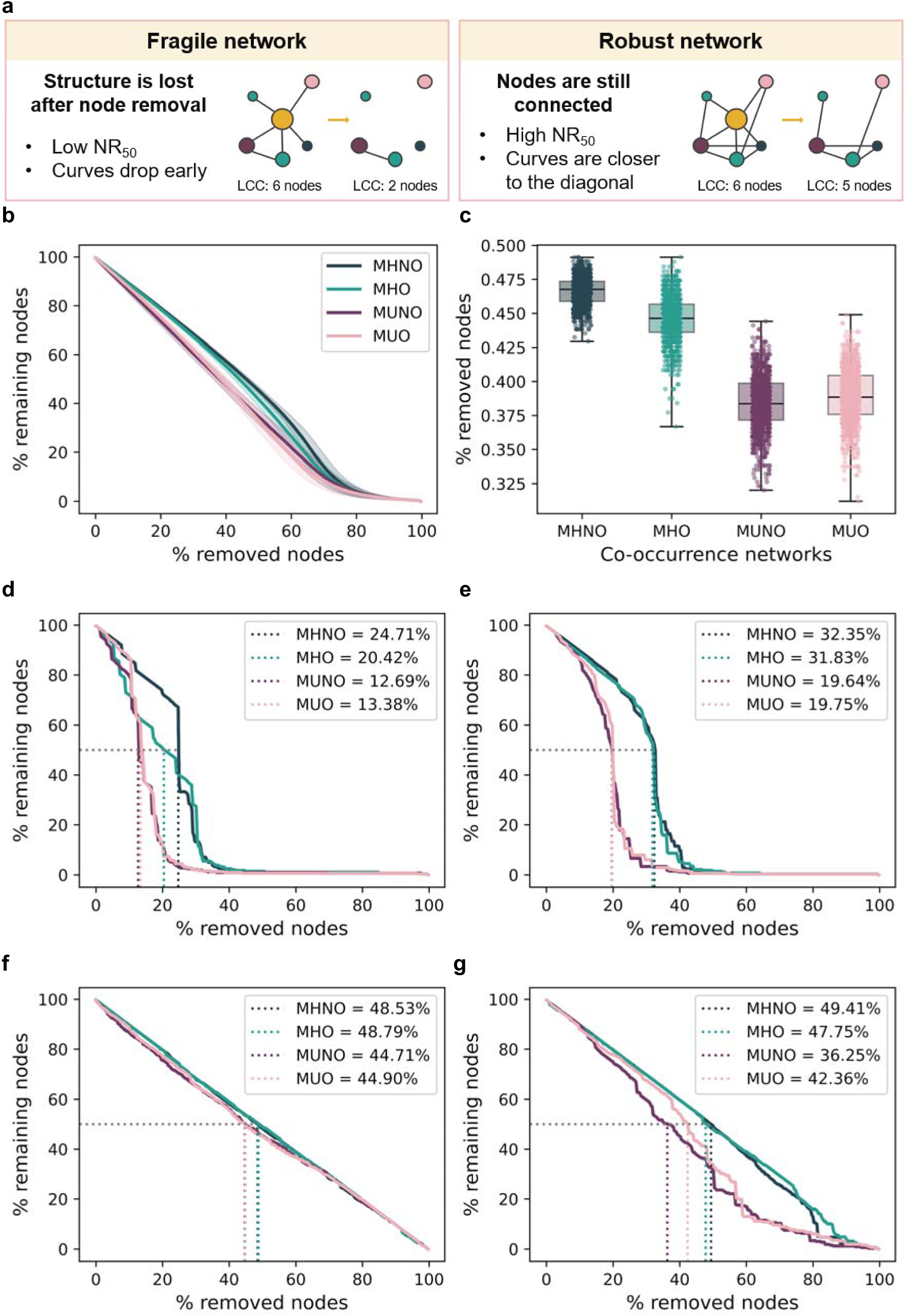
Network stability analysis. **a)** Summary of the approach used to evaluate network stability. **b)** Resistance against random attacks is represented as mean (lines) and standard deviation (shadowed area) after 1000 runs. **c)** NR_50_ values obtained after 1000 runs of random attacks. **d)** Resistance to bottleneck removal. Curves represent the number of nodes in the networks’ remaining LCC compared to the original after sequential removal of nodes with highest betweenness centrality. Legend indicates the percentages of node removal needed to decrease LCC size to 50% of its original value (NR_50_). **e)** Same as (**d**), but assessing robustness against hub removal (i.e, nodes with highest degree). **f)** Network resistance against the loss of taxa based on their decreasing abundance. **g)** Same as **(f)**, but with removal of taxa based on increasing abundance.

First, we performed an exploratory analysis based on random node removal (Figures 5B-C). CNs were highly resistant to these attacks, with LCC orders dropping almost linearly with the number of removed nodes. A difference between MH and MU communities is observed, with MHNO and MHO CNs showing higher NR_50_ (MHNO: 46.59 ± 1.04 %, MHO: 44.64 ± 1.69 %) than their MUNO and MUO counterparts (MUNO: 38.89 ± 2.01 %, MUO: 38.42 ± 2.02 %). Since this approach does not focus on keystone taxa, we expected it to have a small effect on microbial community structure. This might be indicative of how microbial communities respond to subtle perturbations.

Targeted node removal based on betweenness centrality (Figure 5D) results in rapid CN destructuration in MUO and MUNO groups, requiring the removal of only 13.38% and 12.69% of their nodes to disconnect half of the network (NR_50_). The MHNO CN shows the highest robustness, with an NR_50_ of 24.71%, while the MHO CN (NR_50_ = 20.42%) has an intermediate behavior between the MHNO and the MU communities. As for degree-based attacks (Figure 5E), MUO and MUNO CNs behave similarly (NR_50_ = 19.75% and 19.64% respectively), while MHO and MHNO CNs are more resistant (NR_50_ = 31.83% and 32.35%). In this case, no difference between MHO and MHNO graphs is observed. These results show that, while all CNs are sensitive to the loss of keystone taxa, MH communities are more resilient than their MU counterparts, and that the MHO community shows an intermediate resilience between MU and MHNO microbiomes. Network resilience would reflect the ability of the gut microbiota to maintain functionality in stress conditions, such as weight gain, medication intake, or inflammatory or metabolic processes.

We performed a final analysis evaluating network robustness to abundance-based attacks, by removing nodes based on decreasing or increasing mean relative abundances, not necessarily related to their relevance in the community. As shown in Fig. 5F, CNs did not suffer from the loss of the most prevalent taxa, with LCC order decreasing linearly with the number of nodes that were removed. Removal of the less abundant taxa from the CNs had a more powerful effect on their structure, particularly in MU communities (NR_50_ of 36.25% and 42.36% respectively), while in MHNO and MHO CNs, these values increased to 49.41% and 47.75% (Fig. 5G). These results show that MU microbiomes are more heavily dependent on less abundant microbial species than MH communities.

Co-occurrence network approaches provide a new layer of information on how the GM changes in metabolic phenotypes. Our results show that the most prevalent species might not be the most important for community structure and function and that microbial interactions might be more informative of the GM changes underlying certain phenotypes. MHNO microbial communities are characterized by abundant interactions, which are evenly distributed and dominated by SCFA producers. The MHO GM shows small differences in node centrality properties and in network NR_50_ values, which might indicate an underlying change toward metabolically diseased phenotypes. MUNO and MUO microbiomes have fewer connections, dominated by ectopic microbes, that result in unstable microbial communities that are highly sensitive to environmental changes.

## Discussion

Obesity is a complex, heterogeneous condition, requiring tailored treatment strategies as well as public health policies that recognise the diversity in patient phenotypes and needs. Excessive adiposity generates numerous clinical manifestations, forming a continuum where each patient shows a different phenotype. The term “metabolically healthy obesity”, or MHO, was coined with the idea of stratifying patients based on their metabolic health state. Still, there is no agreement on the criteria that characterize MHO, partially due to the heterogeneous nature of disease manifestations related to obesity: some authors focus on inflammatory or insulin sensitivity markers^6^; while others rely on blood pressure, WHR, and the absence of diabetes^87^; or even on the lack of hospitalization for several decades in middle life^88^. Even though these definitions are limited, since they focus exclusively on metabolic health and overlook aspects such as respiratory fitness or joint mobility^2,5^, they are still useful to identify obese patients with better quality of life and a lower cardiovascular or premature mortality risk. Since the GM is involved in several processes related to obesity and metabolic disease, such as immune regulation, glucose homeostasis, or energy metabolism^9^, our work focuses on studying the GM of MHO patients, in particular how obesity and metabolic disease shape microbial interactions and connectivity.

At the anthropometric and biochemical levels, our MHO subjects show intermediate characteristics between MHNO and MUO groups, with smaller waist circumferences, better glycemic and lipidic metabolic state, and lower inflammatory markers than in MUO. This reflects a phenotype in which adipose tissue and glucose metabolism are still functional, although differences in glycemic parameters might be indicating a deteriorated metabolic condition compared to MHNO^1,6^. To further evaluate these patients’ health state as either clinical or preclinical obesity, the function of other organ systems not involved in metabolic regulation should be assessed^2,5^. Despite this limitation, our results reflect the heterogeneity and continuity of obesity phenotypes at the anthropometric, metabolic and inflammatory levels.

Our results from GM exploratory analyses highlight MHNO and MUO as opposite phenotypes, with the largest differences in alpha diversity and differentially abundant species. MUNO and MUO subjects have a less diverse GM^10,12,14^ with higher abundances of pro-inflammatory microbes^10–12^, which are not shared by MHO subjects. These analyses were complemented by the identification of keystone taxa, an alternative approach based on how these microbes interact with each other, rather than on their abundances. This revealed that, while keystone taxa in metabolically healthy communities are predominantly focused on SCFA production, metabolically unhealthy communities feature ectopic microbes among their keystone taxa, a feature that has been described in gut inflammation contexts such as IBD or CRC^83–85^. This highlights a shift in the microbes controlling MH or MU communities at the functional and structural levels. Moreover, MUNO and MUO communities have a weakened interaction network, with changes in connectivity patterns. This results in hampered microbe-microbe communication, along with less stable networks that are highly dependent on keystone species. The higher structural stability of MH communities can reflect a higher capacity to maintain community structure and functionality in the long term, crucial for a healthy GM^89^. If environmental stressors, such as alterations in metabolism, weight gain, or low-grade inflammation, are extended in time, they can alter the structure and function of the microbial community. This would produce a dysbiotic state where cooperation among GM members and plasticity are affected and hence recovery from the loss of microbial species is hindered.

Interestingly, MU communities are also characterized by the increased relevance of microbes with lower relative abundances. In disease contexts, such taxa could provide the necessary characteristics to maintain GM function in the context of metabolic disorders or, conversely, trigger changes in GM structure associated with their onset^90^. These results are further proof that microbial interactions, particularly their stability, are an essential feature to be evaluated in relation to host health, while abundance-based methods can fall short with regard to evaluating GM fitness^18,25,26^.

Current MHO research views this phenotype as an intermediate state between MHNO and MUO^11,16^, and suggests that changes in the GM of MUO subjects are mainly brought on by obesity^10,11^. Thus, we hypothesize that the GM of obese patients with metabolic disorders might be altered in two different ways. On the one hand, obesity would be the main driver of taxonomic changes. On the other hand, the onset of metabolic disease, which can happen independently of excessive fat accumulation, would reshape how microbes interact with each other, affecting their ecological roles and changing the key players in GM structure and function. These alterations are not necessarily reflected in microbial relative abundances, warranting systems biology approaches such as co-occurrence networks to identify and quantify these effects.

Our study population is composed of 959 subjects, representing, to the best of our knowledge, the largest dataset with MHNO, MHO, MUNO, and MUO subjects whose GM has been characterized by shotgun metagenomics. Still, we faced two challenges that limited our MHO sample size: first, many studies focus on phenotypes representing the end of the obesity continuum, looking for comorbidities and complications brought on by excess adiposity, reducing the sample numbers of cases in which these complications are absent. Second, the MHO definition used in this publication requires several biochemical and anthropometric features to be available. Relying on publicly available data has also hampered a more comprehensive evaluation of patient health according to recent guidelines. Finally, our results call for long-term follow-up studies to understand how GM structures may evolve over time, identifying markers of GM stability and proposing potential approaches to prevent the degradation of microbial network structure.

The smaller sample sizes in MHNO and MHO groups have led to an imbalanced dataset, where it was difficult to obtain biomarker information using methods based on microbial abundances, including machine learning approaches. Therefore, we chose a systems biology approach based on co-occurrence networks, which works even in small samples^91^. It should be noted that some limitations, related to the use of samples from different studies or the lack of a gold standard for network-based methodologies in GM studies, might still hinder the interpretability and generalizability of our findings. To mitigate some of these issues, we filtered species with low prevalence, applied batch effect correction methodologies, and adhered to the use of compositionally-aware methodologies. Finally, the interpretation of edge weights, i.e. whether they reflect actual ecological interactions between species, is still under open debate. Indeed, a strict application of network topology analysis to non-physical networks in which edges are inferred by correlation would require an adaptation of these frameworks to consider ensembles of networks and distributions of topological measures. However, an increasing number of studies successfully verifying *in silico*-derived interactions *in vitro*^18–21^ provide support for employing these methodologies.

## Conclusion

Our study aims to describe GM alterations related to different obesity and metabolic phenotypes. To this aim, we have gathered the largest database comprising shotgun metagenomics data from MHNO, MHO, MUNO, and MUO subjects to date and studied their GM through abundance- and network-based approaches. Our findings suggest that, while obesity might be the main driver of GM changes at the taxonomic level, metabolic disorders also drive microbial ecosystem alterations in the human gut. As a result, metabolically disordered GM communities are less biologically stable due to their impaired microbe-microbe interactions, which are dominated by rarer and ectopic microbes. These findings highlight the relevance of network-based approaches to uncover differences in emergent properties, such as community stability, in microbiome research.

## Author Contributions

B.L.-P. Conceptualization, Formal Analysis, Investigation, Methodology, Visualization, Writing – Original Draft Preparation. G.X.B. Data curation. S.R.-T Data curation, Writing – Review & Editing. G.F. Data curation. J.F.-C. Data curation. E.A.-A. Data curation. I.E.-S. Data curation. A.M.-S. Data curation, Writing – Review & Editing. L.P.F. Conceptualization, Writing – Review & Editing. A.R. d M. Conceptualization, Writing – Review & Editing. A.M. Funding Acquisition, Project Administration, Resources, Writing – Review & Editing. R.T. Project Administration, Writing – Review & Editing. J.O.-G. Funding Acquisition, Project Administration, Resources, Writing – Review & Editing. V.P. Conceptualization, Methodology, Supervision, Writing – Review & Editing. L.J.M.Z. Conceptualization, Methodology, Project Administration, Supervision, Writing – Original Draft Preparation. E.C. d S.P. Conceptualization, Funding Acquisition, Project Administration, Resources, Supervision, Writing – Original Draft Preparation.

## Supplementary Material

**Supplementary Figure 1.**
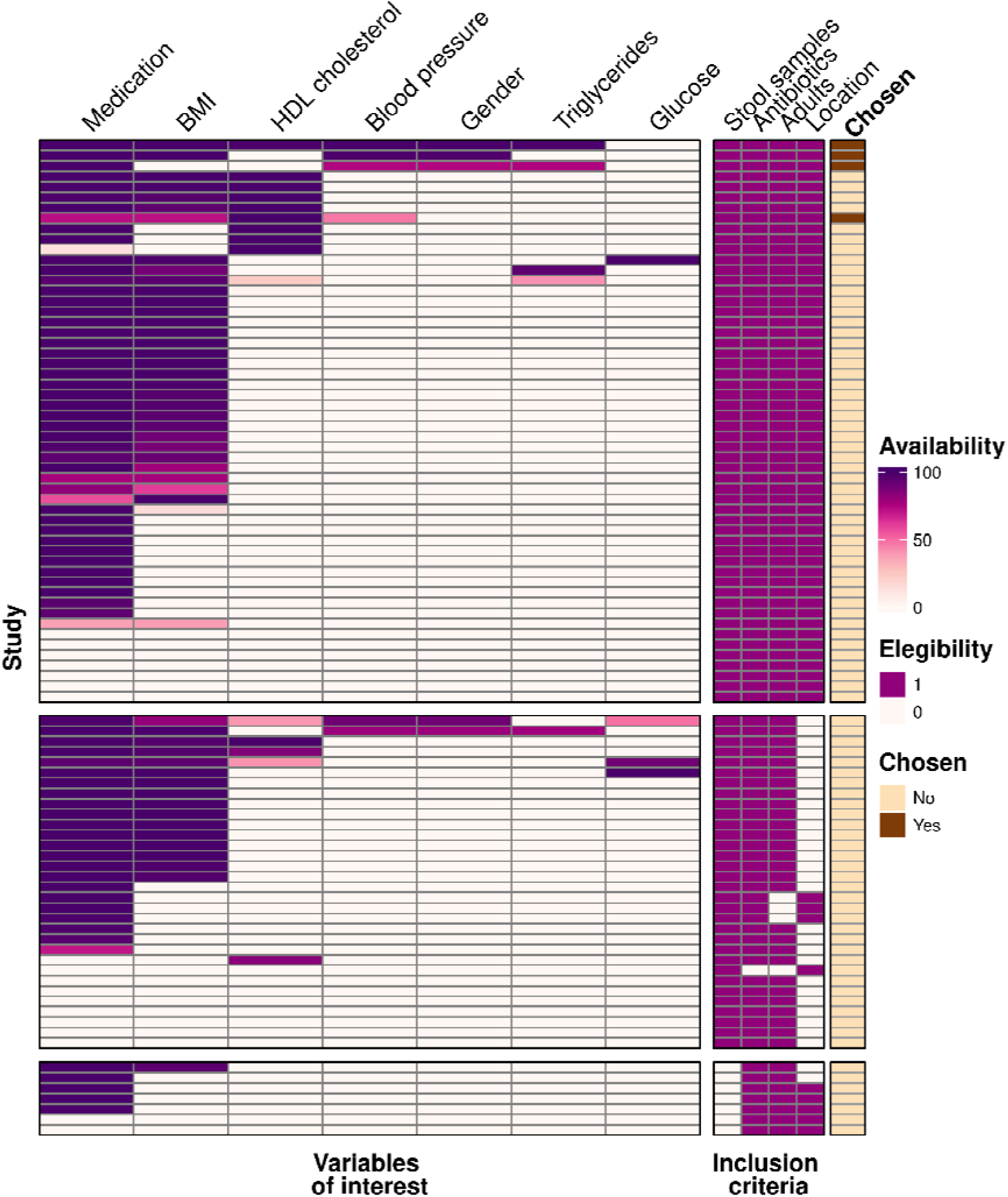
curatedMetagenomicData study search. The left-hand panel shows variable of interest availability in every study from curatedMetagenomicData as the percentage of non-missing values. The right-hand panel shows inclusion criteria met by each study: availability of stool samples, availability of samples from subjects without antibiotic intake, availability of samples from adult subjects, and location. Cells equal to 1 meet these criteria. Rows are separated on three panels based on eligibility criteria fulfillment: the top panel shows studies meeting all criteria, the middle panel shows studies with stool samples that do not meet one of the other 3 criteria, and the bottom panel shows studies without stool samples. Studies chosen for subsequent analyses are indicated in the final column.

**Supplementary Figure 2.**
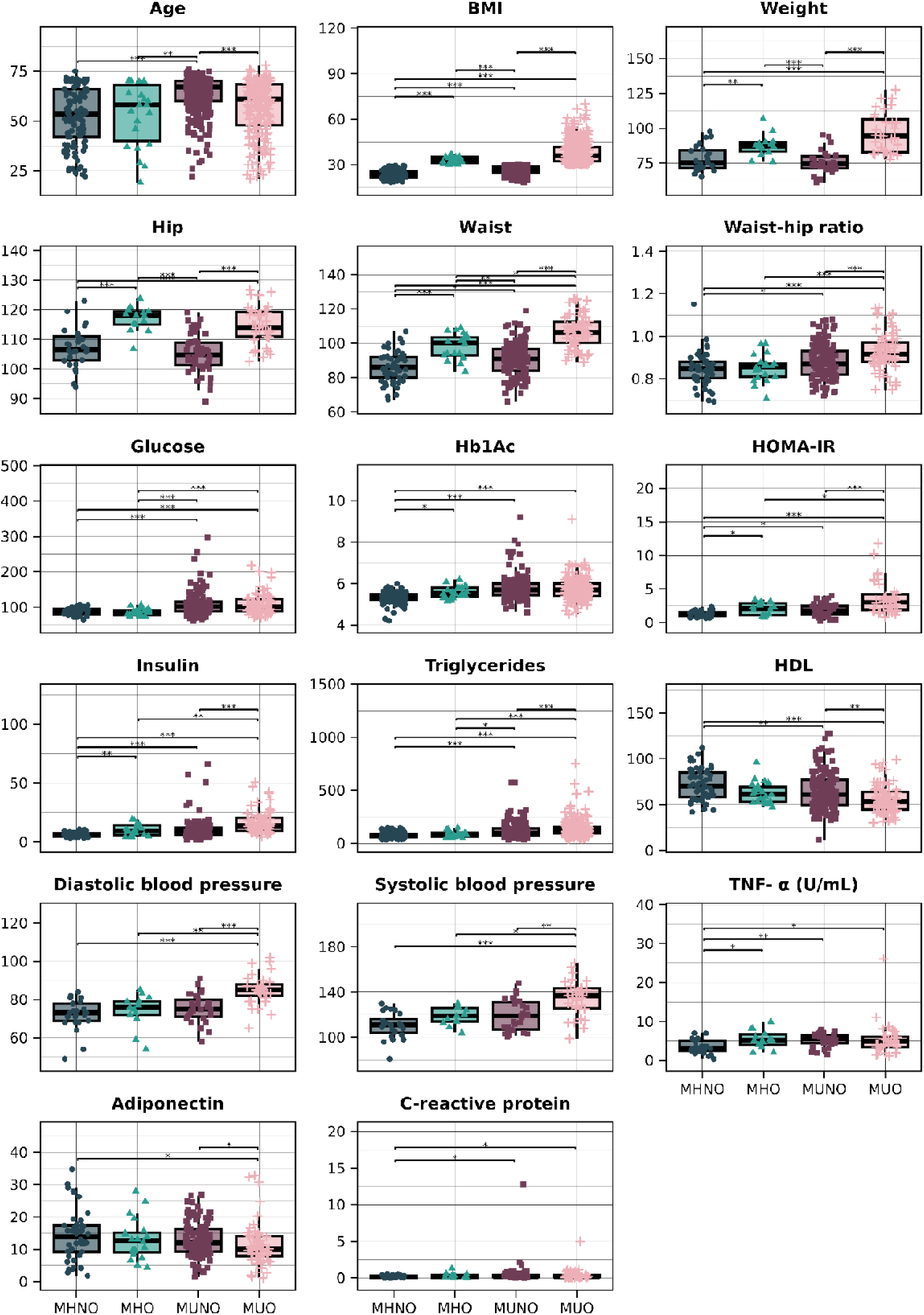
Metadata exploration. Continuous variables with an adjusted p-value < 0.05 in Table 2 were selected. Dunn’s Test for pairwise multiple comparison was used as a post-hoc test. FDR was used to adjust p-values for multiple comparisons.

**Supplementary Table 1. AI4Food patient metadata.**

**Supplementary Table 2. Subject IDs and phenotype classifications obtained from curatedMetagenomicData.**

